# Transcription factor-mediated direct cellular reprogramming yields cell-type specific DNA methylation signature

**DOI:** 10.1101/2023.07.21.549976

**Authors:** Kenichi Horisawa, Shizuka Miura, Hiromitsu Araki, Fumihito Miura, Takashi Ito, Atsushi Suzuki

## Abstract

Direct reprogramming is a technique for inducing the conversion of one type of somatic cell into another by the forced expression of defined transcription factors. Cell differentiation is generally determined by specific gene expression profiles based on distinct genome-wide epigenetic signatures. Although the CpG methylation of genomic DNA is an essential epigenetic factor that affects the transcriptional state of genes, little is known about how DNA methylation changes and what roles it plays in direct reprogramming. Here, we performed comparative genome-wide DNA methylation analyses of mouse embryonic fibroblasts (MEFs) and cells composing organoids formed by intestinal stem cells (ISCs) or induced ISCs (iISCs) that were directly induced from MEFs to investigate the impact of DNA methylation dynamics on direct reprogramming. We found that the methylation state of CpG was similar between cells forming ISC organoids and iISC organoids, while they differed widely from those in MEFs. Moreover, genomic regions that were differentially methylated between ISC organoid- and iISC organoid-forming cells did not significantly affect gene expression. These results demonstrate the accuracy and safety of iISC induction, as they show that the DNA methylation state transitions to a state close to that of ISCs during direct reprogramming from MEFs to iISCs.

## Introduction

A complex molecular mechanism known as epigenetics exists behind the diversity of cells with the same genetic information and enables the readout of specific genetic information on demand. DNA methylation is an epigenetic process in which a methyl group is added at the fifth position of the cytosine pyrimidine ring. Cytosine methylation is classified by sequence context into CG (commonly called CpG), CHH (H = C, A, or T), and CHG sequences, which are regulated by different molecular mechanisms (Sarkies, 2022). CpG sequences are dominant in mammals and are considered major targets of epigenetic regulation (Suzuki and Bird, 2008). In general, CpG methylation near the transcription start site (TSS) has a negative control over gene expression (Deaton and Bird, 2011), whereas CpG methylation can also contribute to the enhancement of transcription (Rauluseviciute et al., 2020; Wan et al., 2015). The DNA methylation patterns of mother cells are accurately transmitted to daughter cells during cell division by a mechanism called maintenance methylation in multicellular organisms, whereas *de novo* methylation, the addition of new methyl groups to specific DNA regions, is another important mechanism for changing the transcriptional state of cells (Greenberg and Bourc’his, 2019). Maintenance and *de novo* methylation are mainly controlled by DNMT1 and DNMT3A/3B, respectively (Chen and Zhang, 2020).

*De novo* methylation is involved in cell differentiation, and its disruption leads to cancer development (Dor and Cedar, 2018). Cellular reprogramming can be artificially induced by using knowledge of cell differentiation and often occurs during cancer development (Horisawa and Suzuki, 2020; Shekhani et al., 2013). Thus, the regulatory system of *de novo* methylation may be an important subject in the study of cellular reprogramming, cell differentiation and cancer development. In fact, the state of DNA methylation dramatically changes not only during cell differentiation but also during cellular reprogramming. Somatic cell nuclear transplantation (SCNT) can reset the state of DNA methylation in transplanted nuclei (Niemann et al., 2008), and the DNA methylation state of induced pluripotent stem cells (iPSCs) transition to a state similar to that of embryonic stem cells (ESCs) during reprogramming (Mikkelsen et al., 2008). However, an inadequate transition of the state of DNA methylation occasionally occurs in SCNT, resulting in a significant reduction in the production ratios of cloned animals (Chan et al., 2012; Zhang et al., 2016). Moreover, during iPSC reprogramming, differentially methylated regions (DMRs) inherited from the original cells (Kim et al., 2010) and appearing ectopically (Lister et al., 2011) are often abnormally generated. Thus, to develop safe and stable cellular reprogramming methods, it is necessary to elucidate, understand, and control the molecular basis of the DNA methylation transition.

As with the technology for inducing iPSCs, direct reprogramming, which can induce a type of somatic cell from another type of somatic cell by the forced expression of defined transcription factors, is considered a promising therapeutic strategy for the treatment of diseases (Horisawa and Suzuki, 2020). Various types of somatic cells, *e.g.* hepatocytes (Huang et al., 2011; Sekiya and Suzuki, 2011), cardiomyocytes (Ieda et al., 2010), and neuronal cells (Vierbuchen et al., 2010), have been generated by inducing direct reprogramming using a combination of transcription factors involved in cell differentiation (Lee et al., 2005; Castro et al., 2011; Hong and Zhang, 2022). A previous study showed that a set of transcription factors used to directly induce neuronal cells synergistically affected the levels of DNA methylation in mouse fibroblasts to establish a neuronal non-CpG methylation pattern (Luo et al., 2019). However, further investigation is required to understand the genome-wide functional role of DNA methylation in direct reprogramming. DNA methylation and demethylation may play essential roles in transcriptional regulation during the progression of direct reprogramming. In fact, it is possible to efficiently induce direct reprogramming by adding 5-aza-2-deoxycytidine, a DNA methyltransferase inhibitor, to the culture medium in combination with the introduction of defined transcription factors into the cells (Koblas et al., 2016). Therefore, it is important to comprehensively understand the role of DNA methylation in direct reprogramming.

In our previous study, we established the method for inducing fetal intestinal progenitor cells (FIPCs) by expressing a set of four genes encoding Hnf4α, Foxa3, Gata6, and Cdx2 in mouse embryonic fibroblasts (MEFs) (Miura and Suzuki, 2017). Under three-dimensional (3D) culture conditions, these induced FIPCs (iFIPCs) formed spherical organoids (SOs) and subsequently gave rise to adult-type induced intestinal stem cells (iISCs) that formed budding organoids (BOs) (Miura and Suzuki, 2017). The morphology and gene expression signature of iISC-derived BOs (iISC-BOs) closely resemble those of intestinal stem cell (ISC)-derived BOs (ISC-BOs), and both iISCs and ISCs can differentiate into functional intestinal epithelial cells, *i.e.* as enterocytes, Paneth cells, goblet cells, and enteroendocrine cells. Moreover, iISCs and ISCs undergo self-renewing cell division and are thus stably maintained in long-term cultures by serial passaging. Transplantation of iISC-BOs and ISC-BOs into a chemically induced colonic injury model results in the reconstitution of intestinal epithelial tissue in damaged colons (Fukuda et al., 2014; Miura and Suzuki, 2017). Based on these findings, we performed a genome-wide and base-resolution methylome analysis using a post-bisulfite adaptor-tagging (PBAT) method (Miura et al., 2012; Miura and Ito, 2018) for MEFs, iISC-BOs, and ISC-BOs, and investigated the transition of the DNA methylation state during direct reprogramming of MEFs to iISCs and the similarities and differences between iISC-BOs and ISC-BOs. Moreover, by combining our present methylome data with our previous transcriptome data (Miura and Suzuki, 2017), we examined the correlation between the abnormal DNA methylation state found in iISCs and the levels of transcription in iISCs.

## Results

### iISC-BOs exhibit a CpG methylation signature that closely resembles ISC-BOs

Genomic DNA was extracted from MEFs, iISC-BOs, and ISC-BOs, and high-resolution methylome analysis was performed using the PBAT method (Figure 1A). The genome-wide methylation states of cytosine-containing sequences such as CpG, CHH, and CHG were compared among the samples. The data showed that CpG methylation signatures were similar among replicates and between iISC-BOs and ISC-BOs, whereas substantial differences were found between iISC-BOs and MEFs, and between ISC-BOs and MEFs (Figure 1B, C). The methylation states of CHH and CHG were highly correlated in comparison to iISC-BOs with their replicates and with ISC-BOs, and MEFs (Figure 1C).

**Figure 1:**
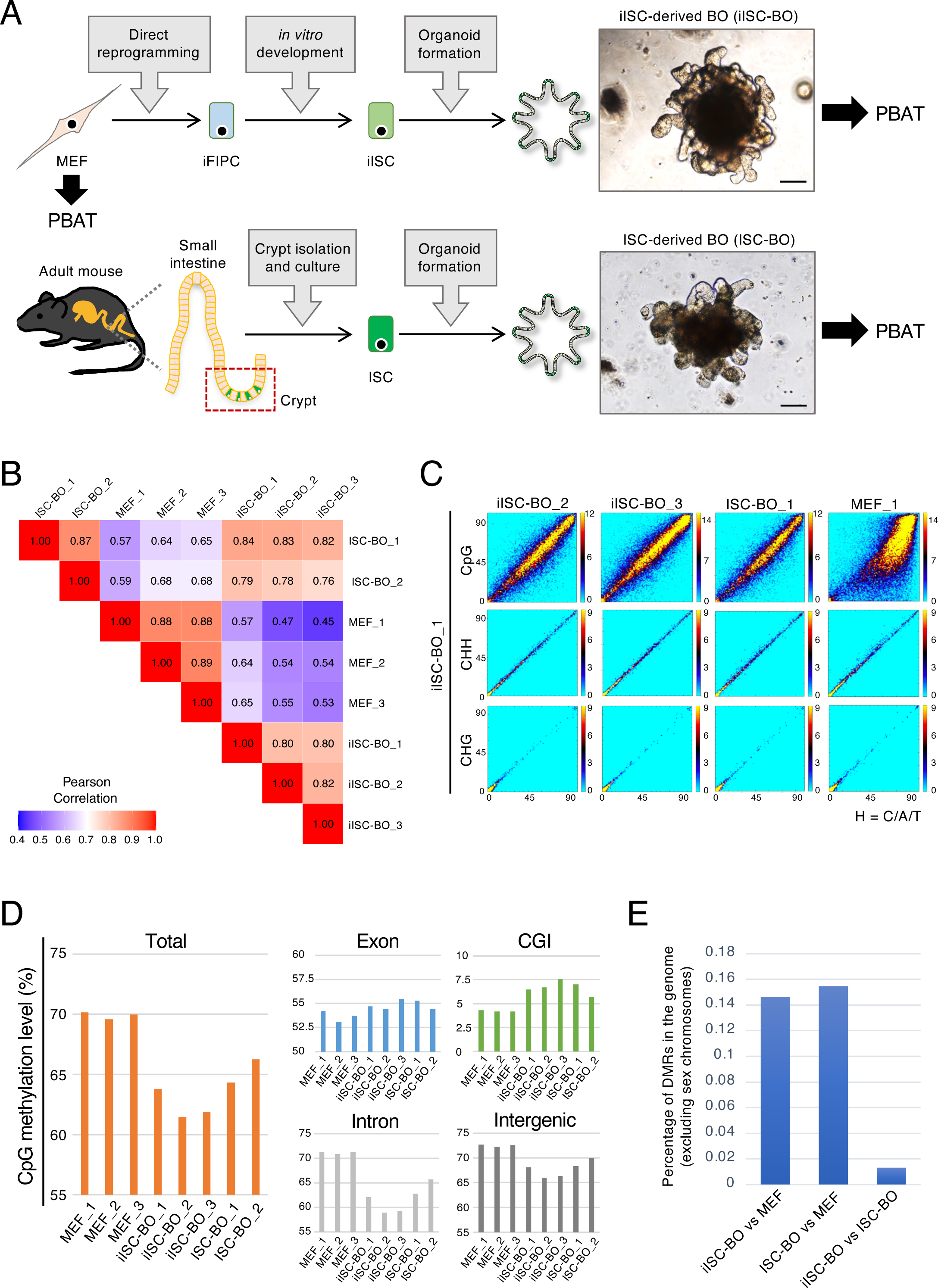
Genome-wide DNA methylation analysis of MEFs, iISC-BOs, and ISC-BOs. (**A**) Scheme of the overall experiment. Photos show bright filed images of BOs. Scale bars, 100 μm. (**B**) Pearson’s correlation analysis of genome-wide CpG methylation between samples. (**C**) Density plots comparing genome-wide CpG, CHH, and CHG methylation status between samples. The window and step sizes were set to be 1 kbp and 500 bp, respectively. (**D**) Averaged CpG methylation levels in each region, which was calculated when the number of reads mapped within a given region was five or more and there were five or more cytosines. (**E**) Genomic occupancy of the DMRs between cells. As a pretreatment of the methylation data analysis, only cytosines assigned with 10 or more reads and located on autosomes were selected.

In a genome-wide view, the total CpG methylation levels of MEFs were 69.6– 70.1%, whereas those of ISC-BOs and iISC-BOs were 64.3–66.3% and 61.5–63.8%, respectively. (Figure 1D, the left panel). The CpG methylation levels of the introns and intergenic regions that occupy most of the genome showed the same tendencies as those of the total regions (Figure 1D, right bottom panels). In contrast, the exons and CpG islands (CGIs) that may have an impact on transcriptional regulation exhibited slightly higher and higher levels of CpG methylation, respectively, in both iISC-BOs and ISC-BOs in comparison to that of MEFs (Figure 1D, right upper panels). Moreover, the percentage of DMRs between iISC-BOs and ISC-BOs was one-tenth that between MEFs and iISC-BOs or ISC-BOs (Figure 1E). Taken together, our data demonstrate that direct reprogramming from MEFs to iISCs is associated with variations in genome-wide CpG methylation and allows the generation of iISC-BOs that have a CpG methylation state similar to ISC-BOs.

### Genome-wide similar distribution patterns of DMRs between MEFs and iISC-BOs or ISC-BOs result in a similar gene expression pattern of iISC-BOs and ISC-BOs

We sought to identify the genomic loci of DMRs between MEFs and iISC-BOs or ISC-BOs to investigate the transition of the CpG methylation state from MEFs to iISCs in more detail. The numbers of DMRs with hypermethylation in iISC-BOs (iISC-high DMRs) or ISC-BOs (ISC-high DMRs) and those with hypomethylation in iISC-BOs (iISC-low DMRs) or ISC-BOs (ISC-low DMRs) were not significantly different, whereas the numbers of iISC/ISC-low DMRs were slightly higher than those of iISC/ISC-high DMRs (Figure 2A). Genome-wide distribution analysis revealed that the genomic loci of iISC-low and ISC-low DMRs and those of iISC-high and ISC-high DMRs were similarly distributed in the intergenic regions and regions of introns and exons, respectively (Figure 2B). To examine how similar distribution patterns of iISC/ISC-low and iISC/ISC-high DMRs affected gene expression, we investigated the expression of DMR-associated genes by reanalyzing our previous transcriptome data obtained from MEFs, iISC-BOs, and ISC-BOs (Miura and Suzuki, 2017). To identify DMR-associated genes, we focused on iISC/ISC-low DMRs and iISC/ISC-high DMRs located within 10 kbp upstream and downstream of the TSS (Figure 2C) because CpG methylation near the TSS affects transcription (Smith and Meissner, 2013). We found that genes associated with iISC-low DMRs and ISC-low DMRs and those associated with iISC-high DMRs and ISC-high DMRs were similarly expressed between iISCs and ISCs compared with MEF (Figure 2D). Thus, the similarity in the genome-wide distribution patterns of iISC-low and ISC-low DMRs and those of iISC-high and ISC-high DMRs may contribute to the similar gene expression patterns of iISC-BOs and ISC-BOs.

**Figure 2:**
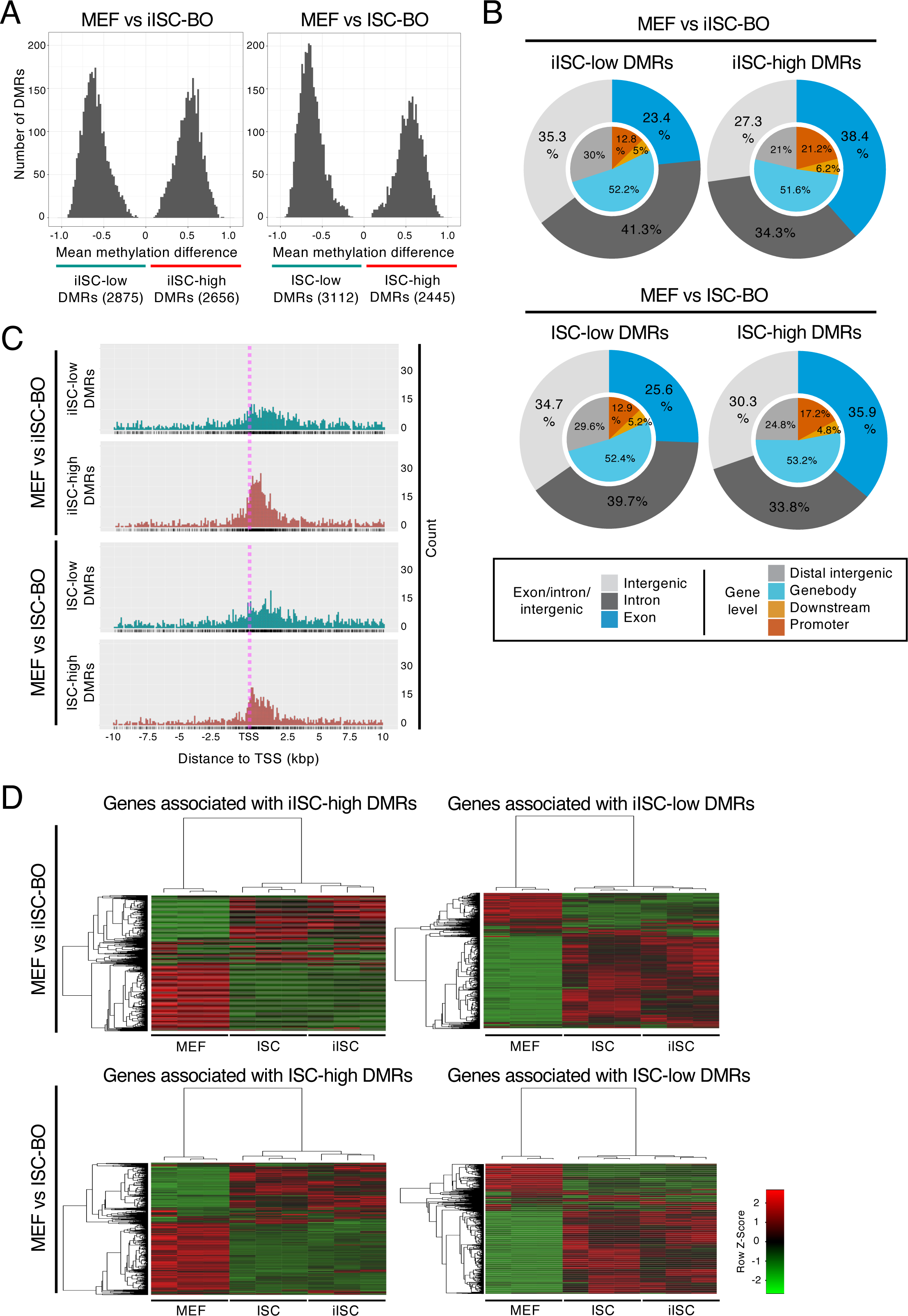
Analysis of DMRs between MEFs and iISC-BOs or ISC-BOs. (**A**) Histograms showing mean methylation difference of DMRs between MEFs and iISC-BOs or ISC-BOs. (**B**) Pie charts showing genomic position of detected DMRs. Outer and inner circles indicate genomic position with different definitions. (**C**) Dot plots showing distribution of DMRs around TSS of DMR-associated genes. (**D**) Heatmaps showing expression level of the DMR-associated genes. As a pretreatment of the methylation data analysis, only cytosines assigned with 10 or more reads and located on autosomes were selected.

### iISC-BOs have hyper- and hypo-methylated DNA, compared with ISC-BOs

As shown above, the CpG methylation state and the resulting gene expression pattern in iISC-BOs were similar to those in ISC-BOs. However, they were not the same. Thus, we examined the genomic loci of DMRs between iISC-BOs and ISC-BOs to explore the differences in CpG methylation states between these two samples. Our data showed that hypermethylation of DMRs in iISC-BOs (hyper DMRs) was more than twice as high as hypomethylation in iISC-BOs (hypo DMRs), suggesting that DNA methylation is induced more frequently than DNA demethylation during direct reprogramming from MEFs to iISCs (Figure 3A). Heatmaps showing the average CpG methylation rates in the hyper-/hypo-DMRs among MEFs, iISC-BOs, and ISC-BOs revealed that these hyper-/hypo-DMRs could be mainly divided into two groups: those that maintain the levels of CpG methylation in the process of direct reprogramming from MEFs to iISCs, and those that represent insufficient or excessive CpG methylation in only iISC-BOs (Figure 3B).

**Figure 3:**
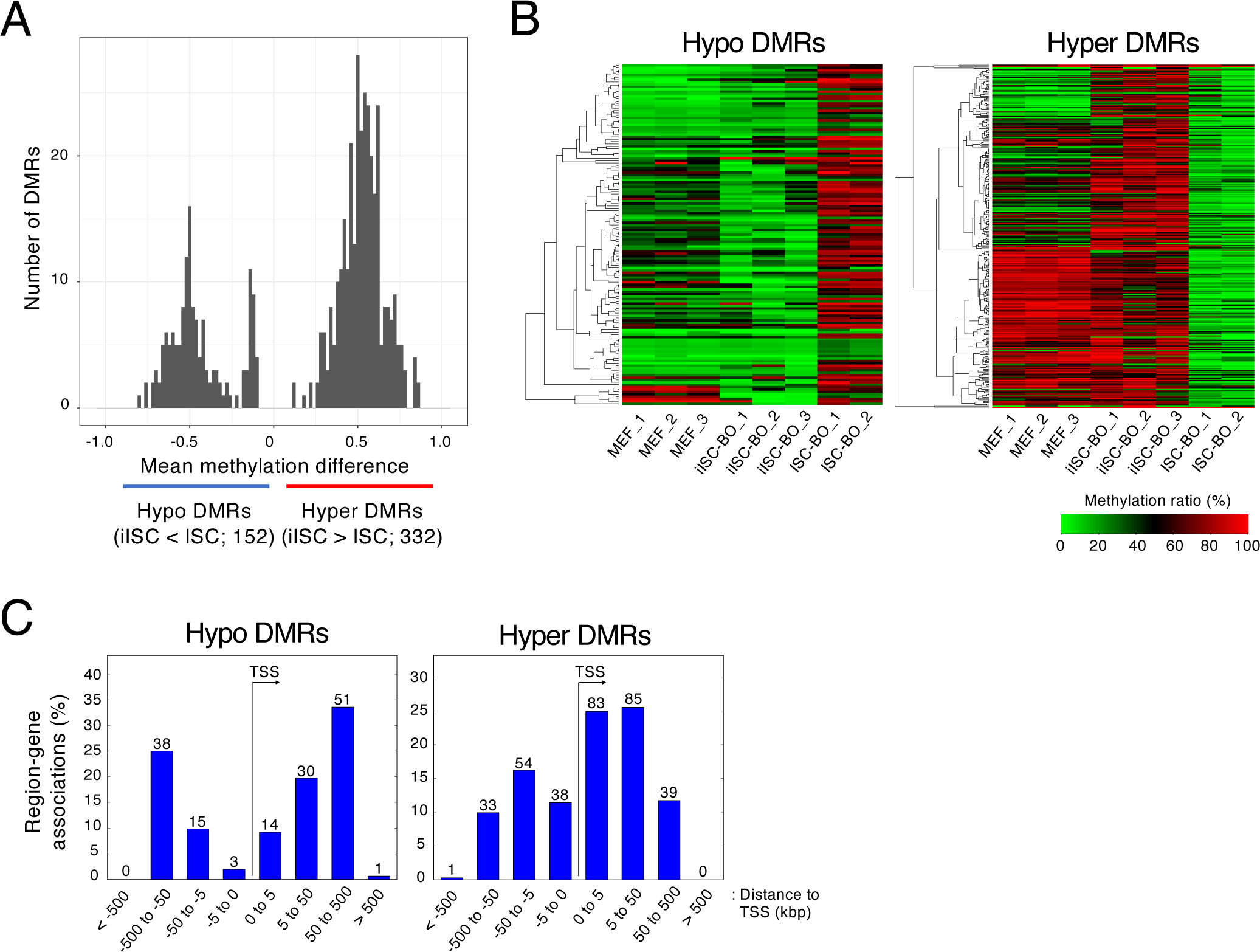
Analysis of DMRs between iISC-BOs and ISC-BOs. (**A**) Histogram showing mean methylation difference of DMRs between iISC-BOs and ISC-BOs. (**B**) Heatmaps showing averaged methylation ratio of detected DMRs in each sample. (**C**) Relative distance of detected DMRs from TSS of DMR-associated genes. The numbers above the bar plots indicate the counts of DMRs. As a pretreatment of the methylation data analysis, only cytosines assigned with 10 or more reads and located on autosomes were selected.

It has been reported that CpG methylation yielded around the TSS affects their transcription (Smith and Meissner, 2013). Thus, we analyzed the genomic loci of the hyper-/hypo-DMRs around the TSS to examine the possibility that these hyper-/hypo-DMRs affect gene transcription. Our data demonstrated that the hyper-DMRs were located in regions distal and proximal to the TSS, whereas the hypo-DMRs were mainly located in regions distal to the TSS (Figure 3C). Thus, hyper DMRs contribute more frequently to the misregulation of gene expression in iISCs than hypo DMRs.

### iISC-BOs have a cellular reprogramming-associated aberrant DNA methylation signature

As shown in Figure 3, an abnormal DNA methylation state may be induced during direct reprogramming. Reprogramming-associated aberrant DNA methylation signatures can be classified into at least two groups. One is an original cell type-specific DNA methylation signature that should be changed but are abnormally maintained in reprogrammed cells (maintained DNA methylation: mDNA methylation), and the other is a DNA methylation signature that is abnormally acquired or erased in only reprogrammed cells (specific DNA methylation: sDNA methylation). To identify regions with mDNA or sDNA methylation in iISC-BOs, DMRs between iISC-BOs and ISC-BOs were divided into two groups based on the overlap with DMRs between MEF and ISC-BOs (Figure 4A). When there was an overlap, iISCs still had an MEF-specific DNA methylation signature, indicating an mDNA methylation signature in iISCs. When there was no overlap, iISCs had a DNA methylation signature that was distinct from that of MEFs and iISCs, indicating an sDNA methylation signature in iISCs. Our data demonstrated that both hyper- and hypo-DMRs could be divided into regions with mDNA or sDNA methylation signatures, and that DMRs with sDNA methylation signatures were found more frequently than those with mDNA methylation signatures (Figure 4B). Genome-wide distribution analyses revealed that both mDNA and sDNA methylation was detected at higher frequencies in exons with hyper-DMRs than in those with hypo-DMRs (Figure 4C). Because hypermethylation of gene body CGIs is considered a result of enhanced transcription (Jeziorska et al., 2017), our data suggest that the mDNA and sDNA methylation signatures found in exons with hypermethylated DMRs reflect errors in transcriptional activation during direct reprogramming from MEFs to iISCs.

**Figure 4:**
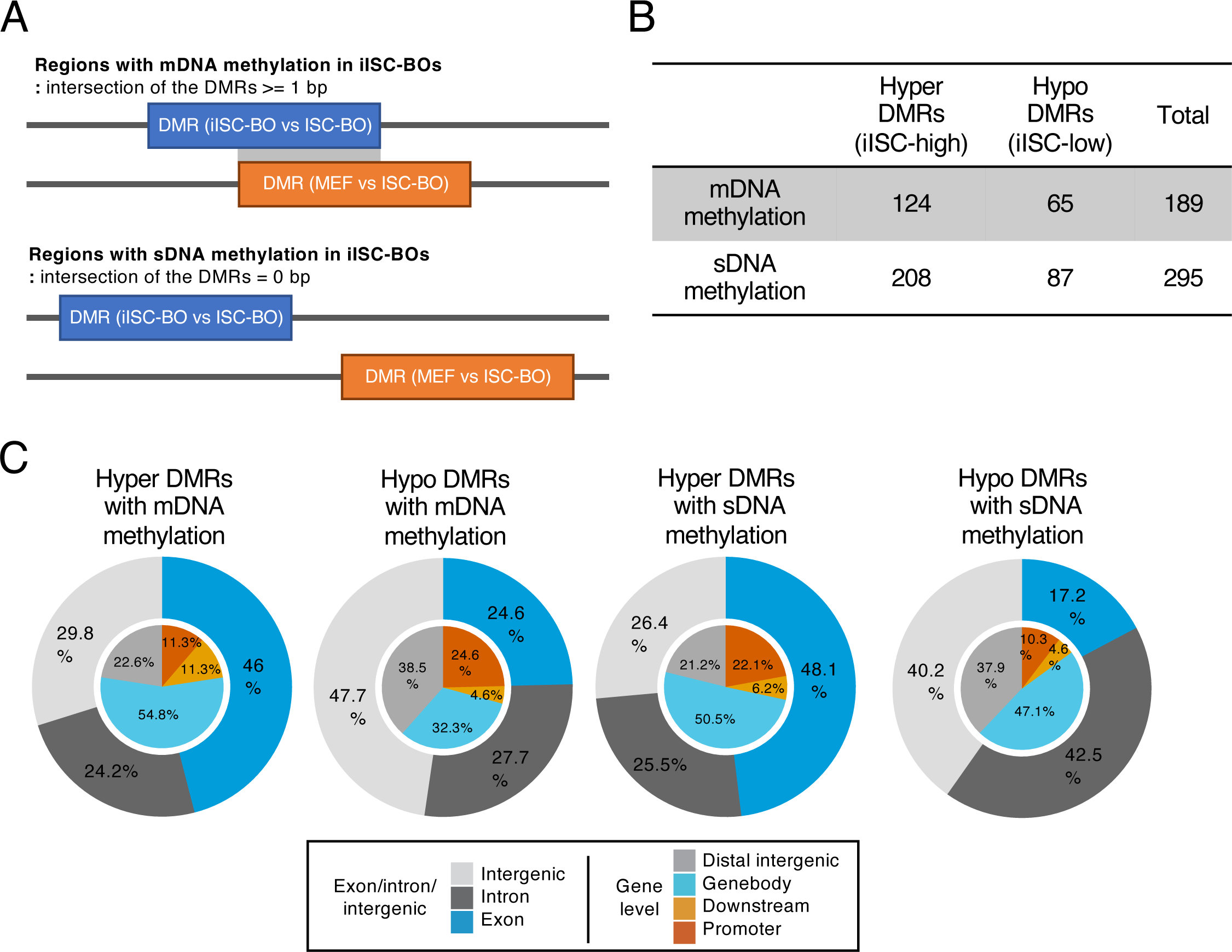
Classification and genomic positioning of abnormally methylated/unmethylated regions in iISC-BOs. (**A**) Definition of DMRs with mDNA and sDNA methylation in iISC-BOs (upper and bottom figures, respectively). (**B**) Number of DMRs in each class: hyper-DMRs with mDNA methylation, hyper-DMRs with sDNA methylation, hypo-DMRs with mDNA methylation, and hypo-DMRs with sDNA methylation. (**C**) Pie charts showing genomic position of DMRs in each class. Outer and inner circles indicate genomic position with different definition.

### Abnormal DNA methylation in iISC-BOs has little effect on gene expression

Our present data suggest that hyper-/hypo-DMRs affect the transcriptional levels of genes and lead to the misregulation of gene expression in iISC-BOs. Our previous study demonstrated that the transcriptional signature of iISC-BOs closely resembles that of ISC-BOs, although a small number of differentially expressed genes (DEGs) were found between iISC-BOs and ISC-BOs (Miura and Suzuki, 2017). Thus, in this study, we examined the relationship between hyper-/hypo-DMRs and DEGs, both of which were identified in iISC-BOs and ISC-BOs. Genes that contained hyper/hypo-DMRs within 10 kb upstream and downstream of the TSS were extracted and divided into DMRs with mDNA or sDNA methylation signatures (Figure 5A). Among these hyper-/hypo-DMR-associated genes, hyper-DMRs were found at high frequencies downstream of the TSS, including the gene body, whereas only a few hypo-DMRs were found within 10 kbp upstream and downstream of the TSS (Figure 5A). In addition, sDNA methylation was found more frequently than mDNA methylation in hyper-DMR-associated genes (Figure 5A). These data suggest that hyper-DMRs are involved in the regulation of DEGs between iISC-BOs and ISC-BOs.

**Figure 5:**
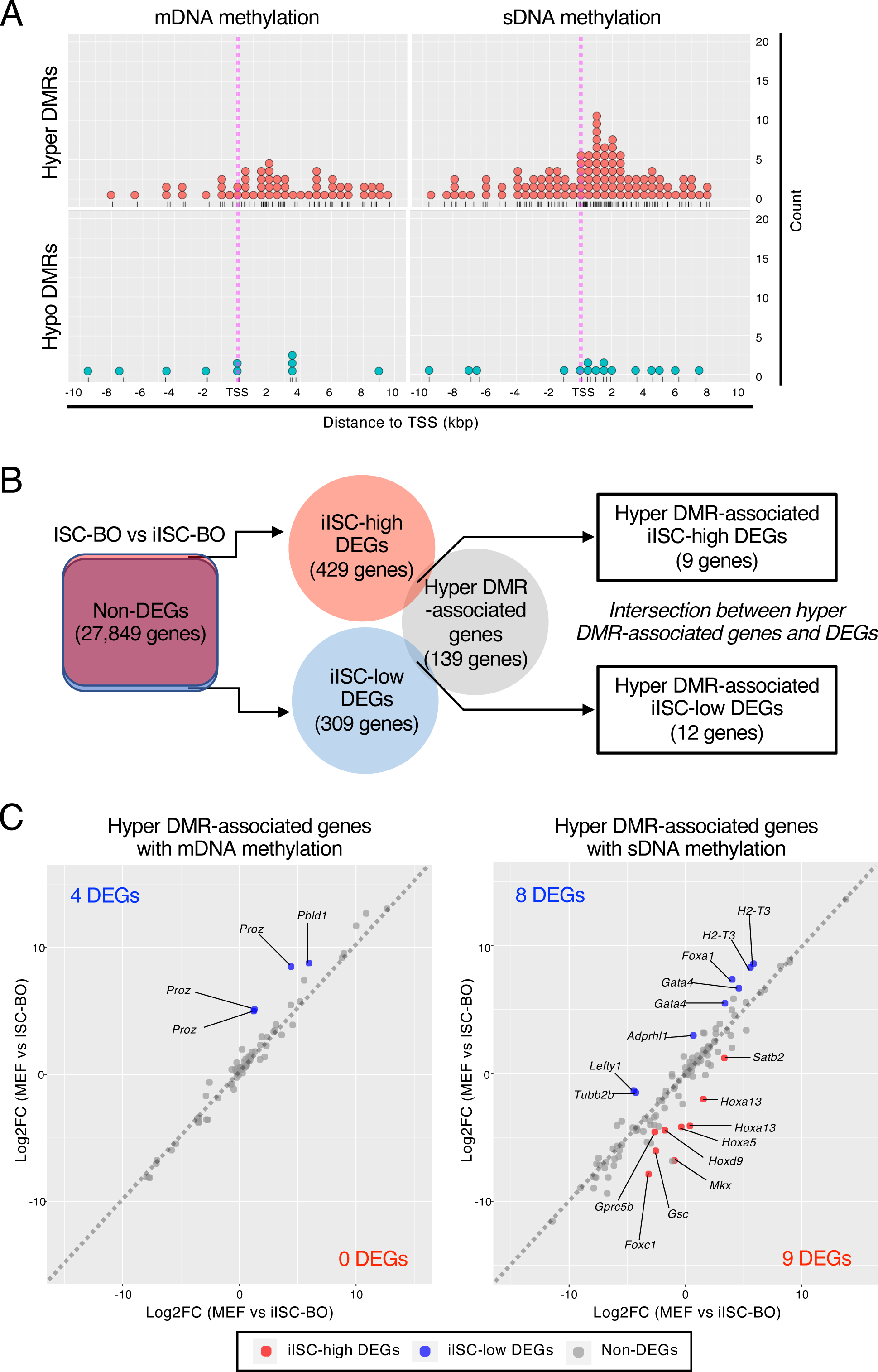
Trans-omic analysis for hyper-DMR-associated genes. (**A**) Dot plots showing distribution of hyper-(upper plots) and hypo-(lower plots) DMRs with mDNA (left plots) and sDNA (right plots) methylation around TSS of DMR-associated genes. (**B**) Right Venn diagram indicates intersection between genes associated with hyper-DMRs and iISC-high or -low DEGs which were detected from a comparative analysis between iISC-BO and ISC-BOs (left Venn diagram). (**C**) Scatter plots showing log-fold change of gene expression (Log2FC) of the hyper-DMR-associated genes between MEFs and ISC-BOs (vertical axis), and these between MEFs and iISC-BOs (horizonal axis). Left and right plots indicate hyper-DMR-associated genes with mDNA and sDNA methylation, respectively.

To examine this possibility, we classified DEGs into genes expressed at higher or lower levels in iISC-BOs than in ISC-BOs, designated iISC-high DEGs or iISC-low DEGs, respectively, and compared each of them with hyper-DMR-associated genes (Figure 5B). Our data demonstrated that the number of genes contained in the hyper-DMR-associated genes and iISC-high DEGs or iISC-low DEGs were 9 and 12, respectively (Figure 5B). These genes comprised only 6.5% and 8.6% of the hyper-DMR-associated genes, respectively. Scatter plots showing fold-differences in the expression of hyper-DMR-associated genes revealed that nine genes have hyper-DMRs with an sDNA methylation signature, while the 12 genes are divided into four and eight genes that have DMRs with an mDNA and sDNA methylation signature, respectively (Figure 5C). Nevertheless, the expression levels of the majority of hyper-DMR-associated iISC-high and iISC-low DEGs (*Foxc1*, *Gprc5b*, *Hoxa5*, *Satb2*, *H2-T3*, *Gata4*, *Lefty1*, *Foxa1*, *Tubb2b*, *Pbld1*, and *Proz*) were significantly upregulated or downregulated in iISC-BOs compared to MEF, in the same manner as in ISC-BOs, while those of other DEGs (*Hoxa13*, *Gsc*, *Hoxd9*, *Mkx*, and *Adprhl1*) did not change significantly (Figure 6).

**Figure 6:**
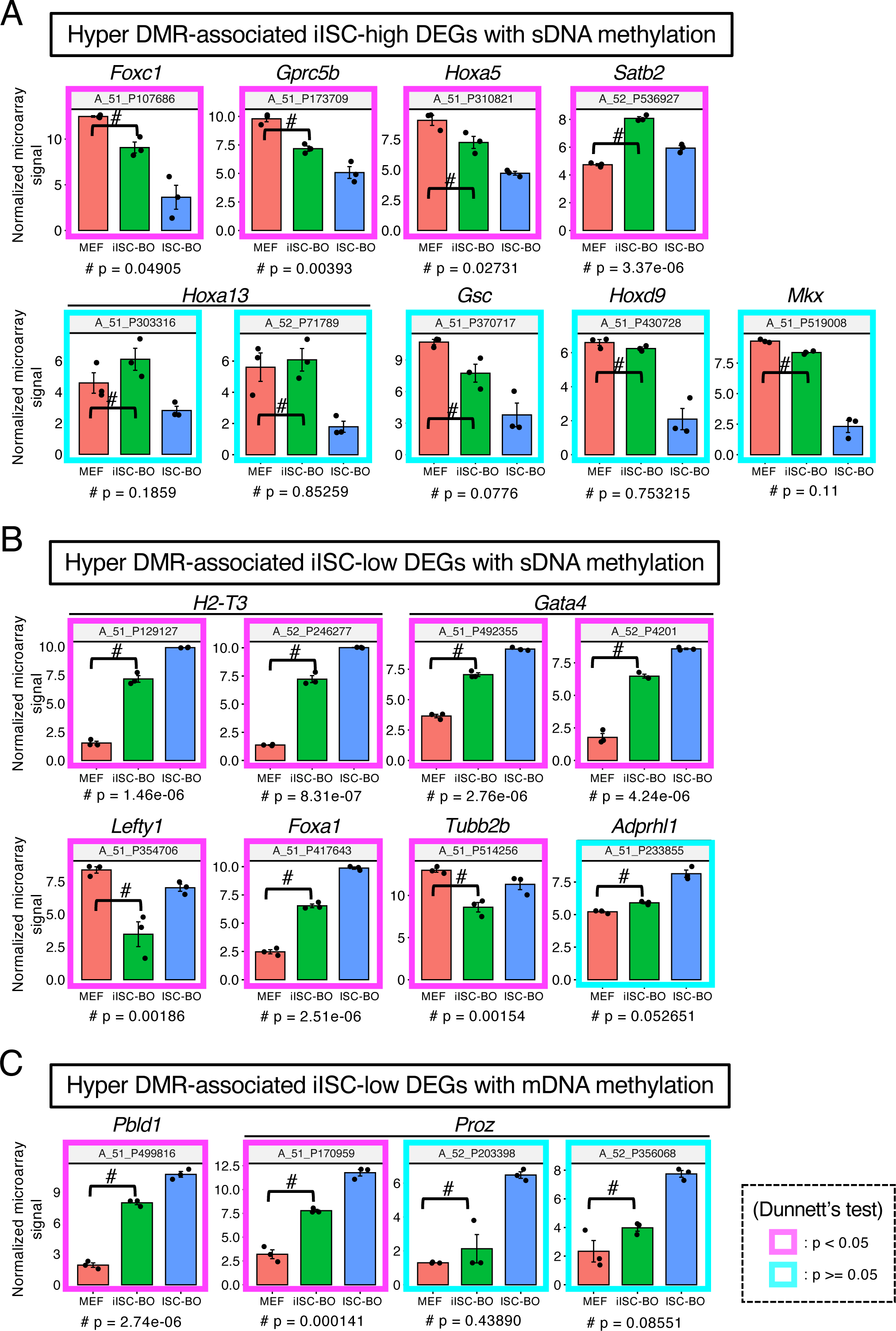
Gene expression analysis for hyper-DMR-associated DEGs. (**A**) Gene expression levels of hyper-DMR-associated iISC-high DEGs with sDNA methylation. (**B**) Expression levels of hyper-DMR-associated iISC-low DEGs with sDNA methylation. (**C**) Expression levels of hyper-DMR-associated iISC-low DEGs with mDNA methylation. Vertical axes of all plots indicate normalized and log-transformed microarray signals. Probe IDs of the microarray corresponding to the transcripts are indicated at the top of each plot. Transcripts showing significant differences in expression between MEFs and iISC-BOs are shown in the magenta box (*p* < 0.05) and those with no difference are shown in cyan boxes (*p* > 0.05). Dunnett’s test was used for statistical analysis.

In addition to the hyper-DMR-associated genes, the hypo-DMR-associated genes also overlapped with iISC-high and iISC-low DEGs, although the number of overlapping genes was only three and five, respectively (Figure 7A). Five genes had hypo-DMRs with an sDNA methylation signature, while three genes were divided into one and two genes that had DMRs with mDNA and sDNA methylation signatures, respectively (Figure 7B). Similar to the case of hyper-DMR-associated iISC-high and iISC-low DEGs, the hypo-DMR-associated iISC-high and iISC-low DEGs (*Foxa1*, *Onecut1*, *Onecut2* and *Ddr2*), except for *Hoxa9*, were expressed at much higher or lower levels than in MEFs and ISC-BOs (Figure 7C). Taken together, our data demonstrated that almost all abnormal DNA methylation states in iISC-BOs may not be involved in the dysregulation of genes that are normally expressed or silenced in ISC-BOs.

**Figure 7:**
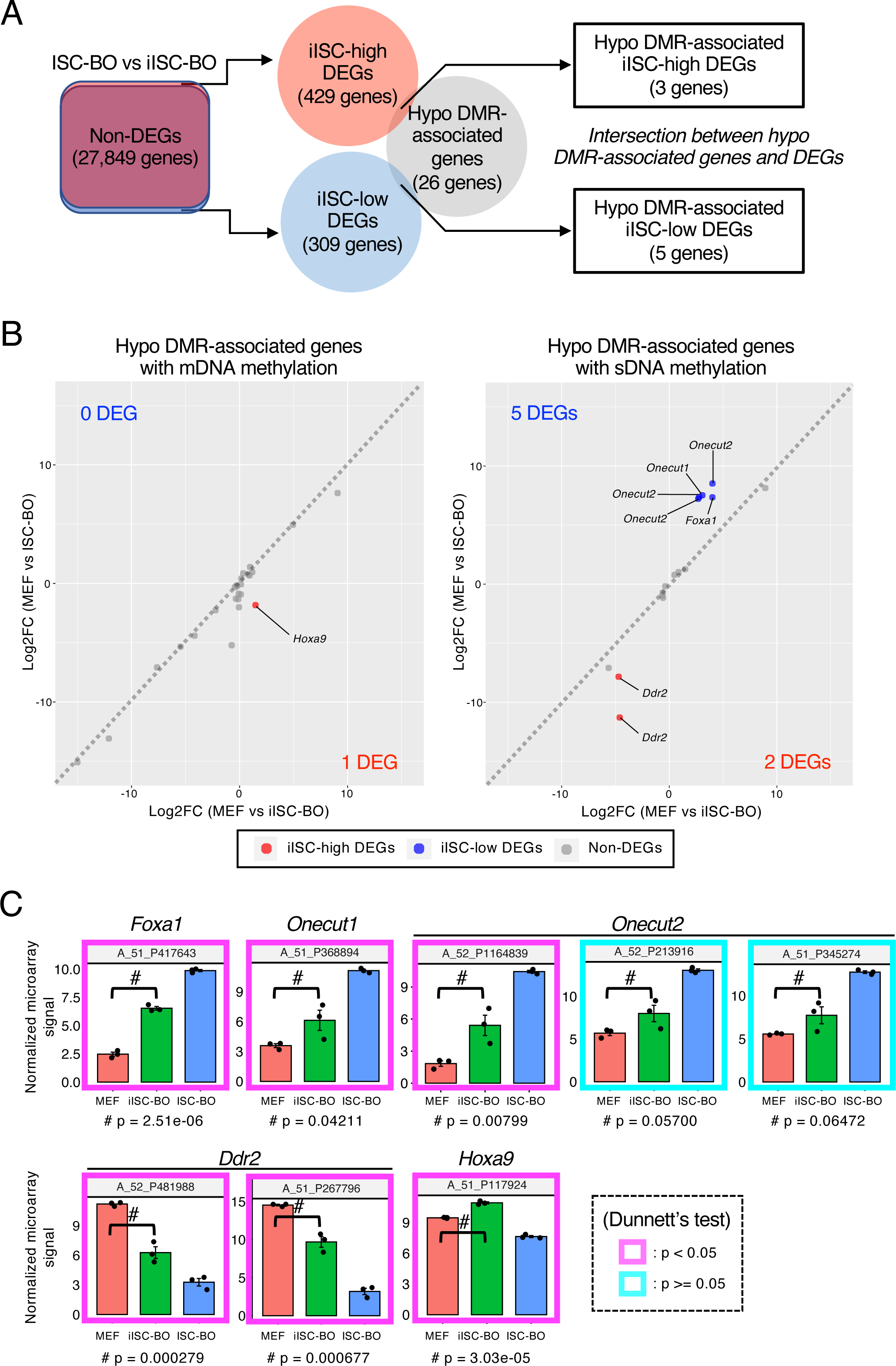
Trans-omic analysis for hypo-DMR-associated genes. (**A**) The right Venn diagram indicates the intersection between genes associated with hypo-DMRs and iISC-high or -low DEGs, which were detected from a comparative analysis between iISC-BO and ISC-BOs (left Venn diagram). (**B**) Scatter plots showing log-fold changes in gene expression (Log2FC) of the hypo-DMR-associated genes between MEFs and ISC-BOs (vertical axis) and between MEFs and iISC-BOs (horizontal axis). Left and right plots indicate hypo-DMR-associated genes associated with mDNA and sDNA methylation, respectively. (**C**) Expression levels of hypo-DMR-associated DEGs. Vertical axes of all plots indicate normalized and log-transformed microarray signals. Probe IDs of the microarray corresponding to the transcripts are indicated at the top of each plot. Transcripts showing significant differences in expression between MEFs and iISC-BOs are shown in the magenta box (*p* < 0.05) and those with no difference are shown in cyan boxes (*p* > 0.05). Dunnett’s test was used for statistical analysis.

## Discussion

Our genome-wide methylome analysis revealed that the methylation state of CpG, but not that of CHH or CHG, changed during the direct reprogramming process of iISC-BOs. CpG methylation, the most frequently observed form of DNA methylation in almost all mammalian somatic cells, is important for transcription (Pelizzola and Ecker, 2011). Previous studies have shown that CHH and CHG methylation may also play important roles in plant and mammalian pluripotency and neuronal cells (Lister et al., 2013, 2009). In the reprogramming of MEFs to iPSCs and neuronal cells, not only the methylation state of CpG, but also that of CHH and CHG changed significantly (Araki et al., 2019; Luo et al., 2019). Thus, it is suggested that variation in CHH and CHG methylation is found in specific cell types and should be corrected during reprogramming from other cell types.

Abnormal CpG methylation signature that affects transcription is often observed during the reprogramming process of iPSCs (Bar and Benvenisty, 2019; Kim et al., 2010). These abnormally methylated regions found in iPSCs are similarly observed at high frequencies in the original cells, which are known as memory DMRs (Bar and Benvenisty, 2019; Kim et al., 2010). These memory DMRs are involved in the differentiation potential of iPSCs (Bar-Nur et al., 2011; Kim et al., 2011, 2010; Lister et al., 2011; Ohi et al., 2011; Roost et al., 2017). Meanwhile, 39% of the abnormally methylated regions found in iISC-BOs were identified as memory DMRs, suggesting that these memory DMRs negatively affect the differentiation potential of iISCs. However, our previous study demonstrated that iISCs can differentiate into functional intestinal epithelial cells in the same manner as ISCs (Miura and Suzuki, 2017). Moreover, among the memory DMR-associated genes, only *Hoxa9* was abnormally expressed in iISC-BOs. Thus, it is suggested that cellular properties and gene expression signatures are less influenced by memory DMRs in reprogramming-induced cells than in iPSCs.

Our present data demonstrated that hyper-/hypo-DMRs within 10 kbp upstream and downstream of the TSS had no significant impact on transcription. However, it is possible that hyper-/hypo-DMRs at a distance of more than 10 kbp from the TSS affect transcription. In particular, hypo-DMRs were predominantly found more than 50 kbp away from the TSS, suggesting that putative enhancers are regulated by DNA demethylation in iISC-BOs. A combined analysis of DMRs with three-dimensional genome structures will allow us to further investigate the effects of DMRs located in regions distal to the TSS on transcription (Gong et al., 2021).

The genome-wide methylation state of iISC-BOs was similar to that of ISC-BOs, although a small number of aberrant methylation sites were observed in iISC-BOs. In iPSCs, the number of aberrant methylation sites gradually decreases because of serial cell division in long-term culture, and the DNA methylation signature of iPSCs approaches that of ESCs (Nishino et al., 2011). Thus, it is suggested that the DNA methylation signature of iISC-BOs gradually resembles that of ISC-BOs by continuous self-renewal cell divisions of iISCs under suitable cell culture conditions. Tissue stem/progenitor cells induced using direct reprogramming technology can obtain a stable DNA methylation state similar to that of tissue-derived allogeneic cells during propagation in culture, which is more suitable for clinical applications than non-proliferative differentiated cells.

## Materials and Methods

### Generation of iISC-BOs and ISC-BOs

Intestinal organoids (iISC-BOs and ISC-BOs) used in this study were prepared as described previously (Miura and Suzuki, 2017). The procedure is briefly described as follows: MEFs were prepared using embryos from E13.5 male and female C57BL/6 mice (Clea, Tokyo, Japan) and cultured in MEF medium (Sekiya and Suzuki, 2011). To induce direct reprogramming, MEFs were infected with retroviruses expressing *Hnf4α*, *Foxa3*, *Gata6*, and *Cdx2*. Adult intestinal crypts were isolated from 10-week-old C57BL/6 male mice (Clea). The reprogramed cells and the cells obtained from adult intestinal crypts were embedded in Matrigel (BD Biosciences, Franklin Lakes, NJ, USA) and cultured in mouse intestinal basal medium with several supplements. The iISC-BOs and ISC-BOs were used after more than 10 passages in 3D culture. All experiments and subsequent analyses were performed according to institutional guidelines and were approved by the ethics committee.

### Whole-genome bisulfite sequencing (WGBS) with a PBAT

Three, two, and three biological replicates were prepared for iISC-BOs, ISC-BOs, and MEFs, respectively, and these samples were individually converted into WGBS libraries. The genomic DNA used for WGBS was prepared using a NucleoSpin Tissue Kit (Macherey-Nagel, Dueren, Germany) according to the manufacturer’s instructions. For sequencing library preparation, 100 ng of purified genomic DNA was spiked with 1 ng of unmethylated lambda DNA (Promega, Madison, WI, USA) and subjected to bisulfite treatment using the EZ DNA methylation gold kit (Zymo Research, Irvine, CA, USA). Library preparation was based on the tPBAT protocol, an improved version of the PBAT strategy (Miura et al., 2012). After library preparation, sequencing was performed using the Illumina HiSeq X Ten system (Macrogen, Tokyo, Japan), assigning one-third of the lanes per sample. The sequenced reads were mapped to the mouse reference genome mm9 combined with the genome sequence of Escherichia phage lambda using BMap as previously described (Miura et al., 2012). The mapped reads were summarized using in-house software, and the basic statistics of the methylome data are provided in Supplementary Table 1. The reads obtained from the biological replicates were treated individually and as groups, depending on the purpose of the analysis. The source codes for the programs used in this study can be downloaded from GitHub (https://github.com/FumihitoMiura/Project-2).

### Transcriptome data analysis

DEGs were detected in the previous study with the GeneSpring software (Agilent, Pal Alto, CA, USA) (Miura and Suzuki, 2017). The log-fold change and normalized signal of each probe were calculated from the raw data. Dunnett’s test was used for statistical analysis.

### Identification of DMRs

The DMRs between MEFs, ISC-BOs, and iISC-BOs were identified using the Metilene program (version 0.2.8) with default parameters (Jühling et al., 2016). DMRs were filtered to retain those containing at least 20 CpGs with a q-value less than 0.05, and a methylation difference larger than 10%.

### Analysis of the genomic location of DMRs

The genomic locations of the DMRs were analyzed using the ChIPpeakAnno program (version 3.28.1) (Zhu, 2013; Zhu et al., 2010) in R (version 4.1.2). The distance between the DMRs and the TSS was determined using the GREAT program (version 4.0.4) (McLean et al., 2010).

## Data availability

Raw WGBS data have been deposited in the NCBI for Biotechnology Information Gene Expression Omnibus (GEO) database (GSE222669). Microarray analysis data for gene expression in MEFs, ISC-BOs, and iISC-BOs are available in the NCBI GEO database (GSE85232).

## Acknowledgments

We thank Drs. Toshio Kitamura, Masafumi Onodera, and Hiroyuki Miyoshi for sharing the Plat-E cells, pGCDNsam and pGCDNsam-ires-EGFP vectors, and the pCMV-VSV-G plasmid, respectively, and Yuuki Honda, Mariko Tasai, Mitsuhiro Kurata, Yoshimi Iwasaki, Emiko Koba, Ryo Ugawa, and Kanako Ichikawa for their excellent technical assistance.

## Author Contributions

K.H. and S.M. performed the experiments, collected samples, and conducted data analyses. H.A., F.M., and T.I. performed methylome analysis using the PBAT method. A.S. contributed to the conception, design, and overall project management. K. H. and A. S. wrote the manuscript.

## Supporting material

**Supplementary Table1: Basic statistics of methylome data**

